# Rapid diversification of the Australian *Amitermes* group during late Cenozoic climate change

**DOI:** 10.1101/2021.04.12.439430

**Authors:** Bastian Heimburger, Leonie Schardt, Alexander Brandt, Stefan Scheu, Tamara R. Hartke

## Abstract

Late Cenozoic climate change led to the progressive aridification of Australia over the past 15 million years. This gradual biome turnover fundamentally changed Australia’s ecosystems, opening new niches and prompting diversification of plants and animals. One example is the Australian *Amitermes* Group (AAG), consisting of the Australian *Amitermes* and affiliated genera. Although it represents the most speciose and diverse higher termite group in Australia, little is known about its evolutionary history. We used ancestral range reconstruction and diversification analyses to illuminate 1) the origin and phylogenetic relationships of the AAG, 2) biogeographical processes leading to the current continent-wide distribution, and 3) timing and pattern of diversification in the context of late Cenozoic climate change. By estimating the first time-calibrated phylogeny, we show that the AAG is a monophyletic group, whose ancestor arrived ~11-10 million years ago from Southeast Asia. Ancestral range reconstruction indicates that Australia’s monsoon region was the launching point for a continental radiation that has been shaped by range expansions and within-area speciation rather than vicariance. We found that multiple arid species diversified from mesic and tropical ancestors in the Plio-Pleistocene, but also observed diversification in the opposite direction. Finally, we show that two pulses of rapid diversification coincided with past climate change during the late Miocene and early Pliocene. Consistent with rapid diversification, species accumulation slowed, likely caused by progressive niche saturation. This study provides a stepping stone for predicting the future response of Australia’s termite fauna in the face of human-mediated climate change.

## Introduction

Australia demonstrates evolutionary challenges that changing climate can pose to organisms and ecosystems. Starting in the late Miocene roughly 10 million years ago (Mya), Australia’s previously warm and wet climate became cool and dry, a period of regional climatic instability known as the “Hill Gap” (Hill, 1994). The widespread rainforests retreated to local refugia (Yeates *et al*., 2002; Cassis *et al*., 2017), while climatic oscillations of the Pliocene and Pleistocene further shaped the present-day arid and semi-arid zones (Byrne et al., 2008). Despite its relatively young age, the Australian arid biome has a rich and unique flora and fauna (Raven & Yeates, 2007; Guzik *et al*., 2011; Powney *et al*., 2010; Ladiges et al., 2011; Andersen, 2016), suggesting either the persistence of ancient lineages or (rapid) radiations in parallel with the increasing aridification of the continent (Crisp *et al*., 2004; Byrne *et al*., 2008). Thus, Australia’s extraordinary biodiversity has not only been shaped by relictualism stemming from the breakup of Gondwana (Barrett & Williams, 1998; Barden & Ware, 2017), but also by vicariance, *in situ* speciation, and phylogeographic structuring of populations in response to local environmental conditions (reviewed in Cassis et al., 2017).

Today, Australia’s arid zone covers roughly 70% of the continent and separates the once predominant mesic biome into eastern and south-western mesic zones (Byrne *et al*., 2008). The mesic biome includes *i.a*. the last remnants of rainforest in Australia and is characterized by high levels of rainfall during the winter season (Byrne *et al*., 2011). Despite receiving similar total amounts of rainfall, the monsoonal tropics in northern Australia are distinguished by summer rainfall, cyclones, and a dry winter season (Bowman *et al*., 2010). While the flora and fauna of each major biome has been extensively studied (Pepper *et al*., 2011; Cardillo *et al*., 2017; Harms *et al*., 2019; reviewed in Byrne *et al*., 2008; 2011; Bowman *et al*., 2010; Rix *et al*., 2015), only a handful of studies have addressed patterns of diversification between Australian biomes at a continent-wide scale (Fujita *et al*., 2010; Owen *et al*., 2017; Brennan & Keogh, 2018), and none so far on Australia’s rich termite fauna (Calaby & Gay, 1959).

Here, we examine the diversification of *Amitermes* Silvestri and allied genera (*Ahamitermes*, *Drepanotermes*, *Incolitermes*, and *Invasitermes*), which we refer to as the Australian *Amitermes* Group (AAG). The AAG forms the most diverse and speciose group of higher termites (Termitidae) in Australia, including about 100 described species (Krishna et al., 2013), many of which play important roles in ecosystem functioning across the continent (Coventry *et al*., 1988; Noble *et al*., 2009, Evans *et al*., 2011). The AAG and other termite lineages are thought to have arrived in Australia relatively recently, 11 to 13 Mya (Bourguignon *et al*., 2017) suggesting that they initially diversified as more or less warm/wet conditions terminated in the late Miocene before facing the challenge of rapidly intensifying aridification in the Plio-Pleistocene. Indeed, diversification in other Australian animal and plant groups coincides with the increase in aridity and the expansion of the arid zone in the last 10 million years (Byrne *et al*., 2008; 2018), including Australian *Coptotermes* and Nasutitermitinae (Lee *et al*., 2015; Arab *et al*., 2017).

AAG species occur across the continent, are adapted to temperate, (sub)tropical, and arid climates, and have persisted in times of severe climate change (Abensperg-Traun & Steven, 1997). This makes them a model system to examine the effects of Australia’s aridification on biogeographical and diversification patterns. Here, we use phylogenetic analyses to see whether this group of higher termites is monophyletic, which would indicate a single introduction event on the Australian continent. We also infer ancestral biomes to examine the impact of the expanding arid zone on species distributions and use diversification analyses to test whether there are changes in the rate of species accumulation coincident with periods of late Cenozoic climate change (*e.g*., during the “Hill Gap” and Plio-Pleistocene).

## Materials and Methods

### Mitochondrial genome sequencing

Sequences were obtained from a total of 87 AAG samples preserved in 70 - 100 % ethanol or RNA*later*, including samples from our own collections (sampled in 2016 and 2019) and the Australian National Insect Collection (ANIC). For collection details, see Supplemental Material, **Tab. S1**.

We used two different sequencing and assembly strategies: (1) long-range PCR followed by deep-amplicon sequencing and (2) ultra-low coverage (1X) whole-genome sequencing (WGS). For the first strategy, whole genomic DNA (gDNA) was extracted from soldier heads using the DNeasy Blood & Tissue extraction kit (Qiagen, Hilden, Germany) following manufacturer’s instructions. Long-range PCRs were performed with PrimeSTAR GXL DNA Polymerases (Takara Bio Europe). Initial sequences generated following Bourguignon *et al*. (2015) were used to develop more effective primers for this group using Primer3 (Untergasser *et al*., 2012) implemented in Geneious R10 ver.10.0.1 (Kearse *et al*., 2012). See Supplemental Material for primer sequences and PCR conditions. The resulting ~8 kb and ~10 kb fragments were mixed in equimolar concentrations and sequenced as paired-end 300 bp reads on an Illumina MiSeq at the Göttingen Genomics Laboratory (G2L, Germany). Amplicon sequencing reads were quality-trimmed using Trimmomatic (Bolger et al., 2014) and assembled with the SPAdes 3.13.0 (Nurk *et al*., 2013) plugin in Geneious Prime using default settings.

For the second strategy, gDNA was extracted from a pool of 4-6 individuals per colony (digestive tracts removed), using the Agencourt DNAdvance Magnetic Bead Kit (Beckman Coulter) or the MagAttract HMW DNA Kit (Qiagen). DNA quality was assessed visually using a 1% agarose gel; quality and concentration were also checked with a Nanodrop 2000 Spectrophotometer (Thermo Fisher Scientific), and a Qubit 2.0 Fluorometer (Thermo Fisher Scientific). Whole genome sequencing (WGS) was performed on a MGISEQ-2000 (Beijing Genomics Institute, BGI, China) resulting in paired-end 150 bp reads. Reads were pre-trimmed by BGI using SOAPnuke1.5.5 (Chen *et al*., 2018).

To retrieve mitochondrial reads from WGS data, we filtered all reads against a customised database of termite whole mitochondrial genome sequences (mitogenomes, **Tab. S1**) using Kraken 2 (Wood *et al*., 2019). Filtered mitochondrial reads were assembled with MitoZ version 2.4 (Meng *et al*., 2019) using the ‘all’ module for paired-end data with default settings (read length: 150 bp; insert size: 300 bp). The resulting mitogenomes were visually checked in Geneious Prime 2020.2.3 (Kearse *et al*., 2012). Unresolved residues and gap characters were manually corrected and the MITOS web server was used for annotation (Bernt *et al*., 2013).

### Mitochondrial data set

The final data set comprised 135 mitochondrial genomes (see Supplementary Material, **Tab. S1**): 87 *Amitermes* and *Drepanotermes* mitogenomes sequenced in this study and 48 sequences from NCBI, including 15 *Amitermes* and *Drepanotermes* mitogenomes and 33 mitogenomes of outgroup taxa to root the phylogenetic inferences.

Control regions of mitogenomes were omitted, as they are generally poorly assembled with short reads due to their highly repetitive character. Each gene was aligned individually using the Muscle algorithm (Edgar, 2004) implemented in Geneious Prime with default settings. Protein-coding genes were aligned as codons. Although, there was no evidence for saturation at third codon positions of protein-coding genes (NumOTU = 32, Iss = 0.265, Iss.cAsym = 0.554) using Xia’s method in DAMBE v.7.2.4 (Xia, 2018), a test for compositional homogeneity using p4 (Foster, 2004; https://github.com/pgfoster/p4-phylogenetics) detected heterogeneity among sequences at the third codon position (chi-square = 1332.07; df = 372; p < 0.0001). Therefore, we excluded those third codon positions from all further analyses resulting in a final concatenated sequence alignment partitioned into four subsets: (a) first, and (b) second codon positions of protein-coding genes; (c) 12S and 16S rRNA genes; and (d) tRNA genes.

### Genetic distances and phylogenetic relationships

Principal Coordinates Analysis (PCoA) was used to visualize genetic (dis)similarity among AAG sequences. Thus, we excluded non-*Amitermes* sequences and *Amitermes* sequences from outside Australia from the final concatenated sequence alignment. SNPs were called with the R package ‘adegenet’ ver. 2.1.1 (Jombart, 2008; Jombart & Ahmed, 2011) using R version 3.6.3 (R Core Team, 2020). The PCoA was performed in R with vegan ver. 2.5-6 (Oksanen *et al*., 2019).

The final concatenated sequence alignment was analyzed in a maximum likelihood framework using IQ-TREE ver. 2.0.6 (Minh *et al*., 2020). We used the implemented algorithm of PartitionFinder (Lanfear *et al*., 2012) to search for the best-fitting partitioning scheme, which increases model fit by reducing overparameterization. The model selection procedure was immediately followed by tree reconstruction (‘-MFP+MERGE’ command). We performed 1000 replicates each for both ultrafast bootstraps (ufBS) (‘-bb’ command) and SH-aLRT tests (‘-alrt’ command). Nodes were classified as “robust”, if recovered support values for SH-aLRT/ufBS were ≥ 75 %.

To date our phylogeny, we used four termite fossils as internal calibrations (see **Tab. S2**): (1) †*Nanotermes isaacae*, (2) †*Reticulitermes antiquus*, (3) †*Microcerotermes insularis* and (4) †*Amitermes lucidus*. †*Nanotermes isaacae* is the oldest known Termitidae fossil and at least 47.8 million years old (Engel *et al*., 2011); we used it to calibrate the Termitidae + sister group. †*Reticulitermes antiquus* is known from Baltic amber (Engel et al., 2007) (minimum age of 33.9 million years) and likely represents a stem group *Reticulitermes* according to Bucek *et al*. (2019), so we used it to calibrate *Reticulitermes* + sister group. (3) †*Microcerotermes insularis* and (4) †*Amitermes lucidus* are both known from Dominican amber (minimum age of 13.8 million years) and their generic assignment is clearly established (Krishna & Grimaldi, 2009). The fossil calibrations were implemented as exponential priors using the aforementioned minimum fossil ages as reported on the Paleobiology Database (www.paleobiodb.org; accessed 30 July 2020) as minimum age constraints. Soft maximum bonds (97.5 % probability) were deliberately chosen to be very old (**Tab. S2**), because the fossil record of termites is highly fragmentary, which can lead to underestimated node ages (Ho & Philips, 2009).

For each partition of the final concatenated sequence alignment, we used the bModelTest package (Bouckaert & Drummond, 2017) to average over all transition/transversion split models (31 different models in total; see Bouckaert & Drummond, 2017) implemented in BEAST 2.6.1 (Bouckaert *et al*., 2019) using reversible jump Markov Chain Monte Carlo (rjMCMC). This approach takes into account uncertainty in the model selection process and subsequent bias in estimates based on single models (Bouckaert *et al*., 2017). In all analyses, we used an uncorrelated lognormal relaxed clock model of rate variation across branches (Drummond *et al*., 2006) and a birth-death speciation process as tree prior.

We performed three independent MCMC runs, sampling tree and parameter values every 25,000 steps over a total of 500 million generations. Convergence and effective sample sizes were checked with Tracer v1.7.1. (Rambaut *et al*., 2018), with a quarter of samples removed as burn-in. MCMC runs were combined with LogCombiner v2.6.2 (Drummond & Rambaut, 2007) and a maximum-clade-credibility tree was obtained using TreeAnnotator v2.6.0.

### Biogeographic history

We used our time-calibrated tree to compare biogeographic models and estimate ancestral ranges using the maximum-likelihood approach implemented in the R package BioGeoBEARS v.1.1.2 (Matzke, 2013). We compared three different biogeographic models implemented in BioGeoBEARS: DEC, DIVALIKE, and BAYAREALIKE. These biogeographic models include different cladogenetic processes: (1) DEC (Dispersal-Extinction-Cladogenesis) includes subset sympatry; (2) DIVALIKE (likelihood interpretation of DIVA) includes widespread vicariance; and (3) BAYAREALIKE (likelihood interpretation of BayArea) includes widespread sympatry. The best-fitting model was assessed with the Akaike Information Criterion (AIC; Akaike, 1974).

Prior to analysis, we excluded all outgroup taxa and non-Australian *Amitermes*. Additionally, we pruned the tree of splits younger than 1.5 million years, retaining only a single representative of each species/independent evolutionary unit to avoid spurious “speciation” events as recommended on the BioGeoBEARS website (http://phylo.wikidot.com/biogeobears-mistakes-to-avoid#toc6). This resulted in a tree with 72 terminal tips, which were assigned to the four major biomes in Australia (modified from Fujita *et al*., 2010; abbreviated as follows: S, mesic south-western zone; A, arid zone; M, monsoonal tropics; and E, mesic eastern zone) depending on the collection site (**Fig. 1c** and **Tab. S3**). Where tips could be unambiguously attributed to a described species, occurrence records from the Atlas of Living Australia (http://www.ala.org.au. Accessed 10 July 2020) were used to assist biome assignment. We allowed a maximum of two biomes to form a species range, while excluding all combinations of non-adjacent biomes (*i.e*., “S+M” and “S+E”), which resulted in 9 possible ranges in total.

**Figure 1:**
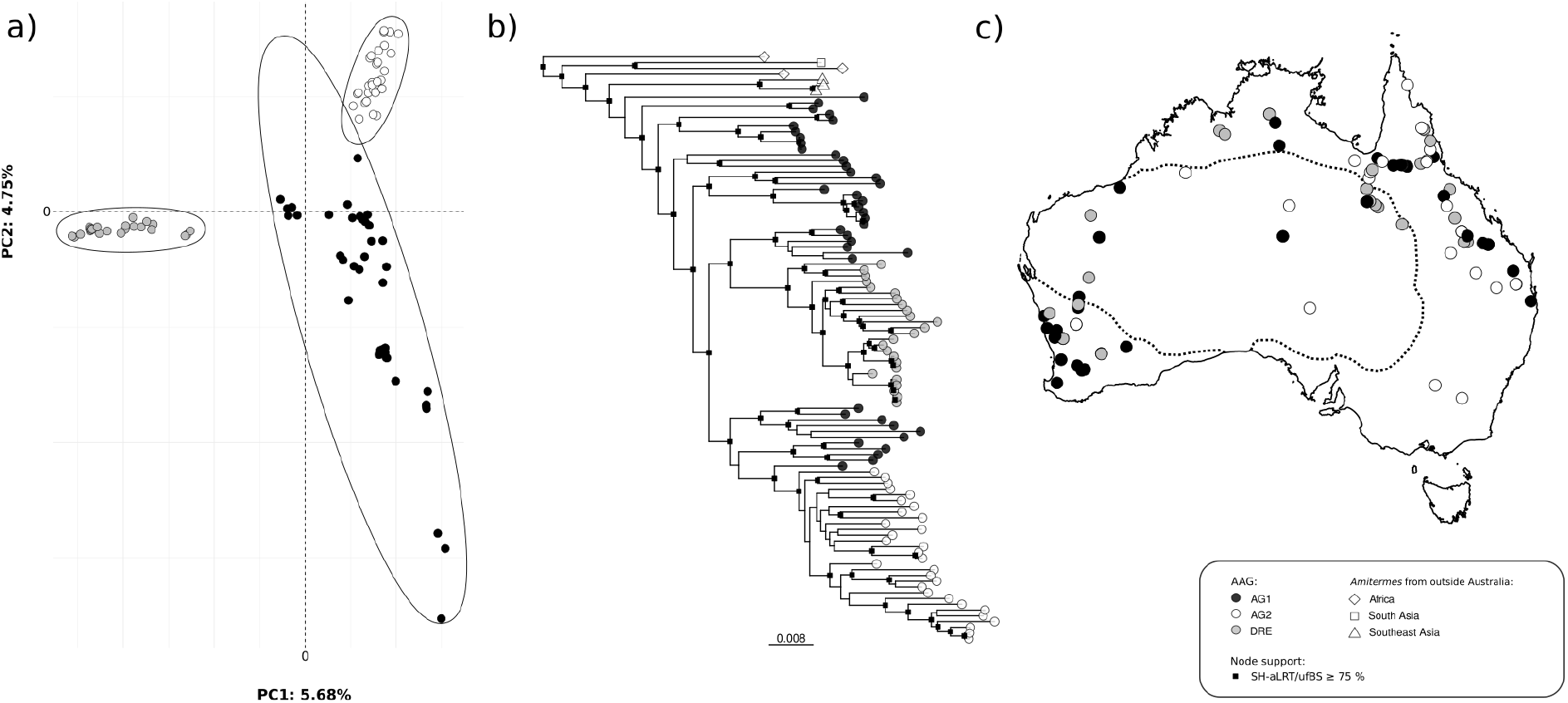
a) The PCoA showed three distinct genetic clusters within the AAG, b) two of which (DRE and AG2) are recovered as monophyletic by the maximum likelihood analysis. c) Map shows sampling locations of AAG taxa included in this study in relation to the arid zone (dashed line; modified from Fujita *et al*., 2010).

We used 100 Biogeographical Stochastic Mappings (BSMs) to obtain the overall probabilities of the anagenetic and cladogenetic events (Matzke, 2016; Dupin *et al*., 2017), which depend on the geographic distributions, the time-calibrated phylogeny, and the best-fitting model. This allowed us to quantify the relative role of dispersal and vicariance at cladogenesis in the diversification of the AAG.

### Model and rates of diversification

The temporal pattern of lineage diversification was visually assessed with a semi log-scaled Lineage-through-time (LTT) plot in the R package phytools (Revell, 2012) using the pruned time-calibrated tree (see above), as well as 500 simulated LTTs assuming a pure-birth process of the same duration and resulting in the same total number of species. The γ statistic (Pybus & Harvey, 2000) was simultaneously calculated, which can detect whether the net diversification rate deviated over time from a pure birth model (standard normal distribution with a mean of 0). We conducted a Monte Carlo constant rates test (MCCR test; Pybus & Harvey, 2000) implemented in the R package LASER (Rabosky, 2006), to account for incomplete sampling (Fordyce, 2010). The test mimicked incomplete sampling by randomly pruning taxa from phylogenies, which were simulated to the full size of the group (i.e., about 100 described species according to Krishna *et al*. (2013)). We used 10,000 replicates under the null hypothesis of a constant rate pure-birth diversification process.

In addition to the MCCR test, we compared the fitting of alternative evolutionary models in respect to our LTT plot using the run_diversification_analyses.R script (Condamine et al., 2018). This analytical pipeline includes the R packages RPANDA (Morlon *et al*., 2016) and DDD (Etienne *et al*., 2012). Both approaches take into account the absence of species (both extinct and missing from the phylogeny) and how this is likely to affect historical diversification rates. We used four different models, of which two assume constant diversification rates including the pure-Birth (or Yule) model assuming no extinction, and the constant rate birth-death model (CR) with extinction but constant rates of speciation and extinction over time and among lineages. The other two models assume diversity-dependence, namely the density-dependent linear (DDL+E) and the density-dependent exponential (DDX+E) models. Both models quantify diversification rates as functions of changes in species accumulation over time. The DDL+E model assumes a linear dependence of speciation rate with extinction (E), while the DDX+E model assumes an exponential dependence of speciation rate with extinction.

We fitted all models under three alternative scenarios to account for different proportions of missing species in our phylogeny. Although the number of species currently reported for the AAG is comparatively high, the ‘true’ diversity is likely much higher. Hence, we assumed that the group (a) consists of the currently reported number of species (100), (b) includes more species (150) than known today, and (c) is actually much larger than currently described (250). In other words, we assumed that the sampling fractions in our phylogeny were 72%, 48%, and 29%, respectively. The AIC approach was used to evaluate the best-fitting model, reported as the bias-corrected AIC version (AICc; Burnham & Anderson, 2002; Posada & Buckley, 2004). To determine the goodness of fit of candidate evolutionary models, we used the lowest AICc score and ΔAICc scores, where the differences between the lowest (or best) AICc and the AICc of each alternative model are calculated, and thus, the best model has ΔAICc = 0.

To complement the model-fitting approach, we employed BAMM 2.5.0 (Bayesian Analysis of Macroevolutionary Mixtures; Rabosky et al., 2013; Rabosky, 2014; Rabosky *et al*., 2014; bamm-project.org), which can automatically detect diversity-dependence on phylogenetic trees, shifts in diversification rate through time, and key innovations. We ran BAMM for 100 million generations with default parameters and a burn-in of 25% using the same pruned time-calibrated tree as above. To account for incomplete taxon sampling, we implemented 0.72 as sampling fraction. Since prior settings can have a substantial impact on BAMM analyses (Moore *et al*., 2016), we used a gradient of values for the prior on the number of shifts in diversification (compound Poisson process), ranging from the default value of 1.0 (higher probability of no rate shift) to 0.1 (higher probability of multiple rate shifts), with a step of 0.1. The best-fitting run (*i.e*., highest posterior probability for the number of shifts) was selected for downstream analyses. The R-package BAMMtools 2.1 (Rabosky *et al*., 2014) was used to check for convergence (ESS > 200) and mixing of the MCMC chain in each analysis.

## Results

### Genetic distances and phylogenetic relationships

Phylogenetic reconstructions based on Bayesian and ML inferences consistently recovered a monophyletic AAG (> 85 % SH-aLRT/ufBS values and posterior probabilities), which is sister to *A. dentatus* from Southeast Asia (**Fig. 1**). The PCoA indicated three major groups within the AAG (**Fig. 1**): *Amitermes* group 1 (AG1), *Amitermes* group 2 (AG2), and *Drepanotermes* (DRE). The latter two formed independent monophyletic crown groups nested within the paraphyletic group AG1, which were well-supported in all phylogenetic analyses (> 95 % SH-aLRT/ufBS values and posterior probabilities; **Figs. 1** and **S1 & 2**).

Divergence estimates, based on the model-averaging approach (bmodeltest) without third codon positions, dated the split between the AAG and the Southeast Asian species at 10.99 Mya (95% HDP: 9.52-12.81 Mya) (**Fig. S2**). Two major divergence events, *Drepanotermes* at 6.5 Mya (95% HDP: 5.6-7.67 Mya) and AG2 at 5.55 Mya (95% HDP: 4.77-6.45 Mya), correspond roughly to the end of the paleoclimatic ‘Hill Gap’ (10-6 Mya) in the late Miocene (**Fig. S2**). An estimated 12% of DRE and 53% of AG2 lineages diverged during the Pliocene (~5.3-2.6 Mya) compared to 33% of AG1 lineages, while recovered divergence time for 35 out of 95 lineages within the AAG were < 1.5 Mya (**Fig. S2**).

### Biogeographic history

The AIC model selection favored the DIVALIKE model, which was 1.8 AIC units lower than DEC and 29.1 AIC units lower than BAYAREALIKE (**Tab. 1**). The models that included vicariant processes (DEC and DIVALIKE) gave very similar histories compared to the BAYAREALIKE model (results under DEC and BAYAREALIKE are available in **Fig. S3**). We expected that vicariant speciation played a substantive role in the evolution of the AAG, because it has shaped the evolutionary trajectory of many arid zone taxa (Cracraft, 1982; Crisp & Cook, 2007; Rabosky *et al*., 2007).

**Table 1:** Summary statistics of three biogeographic models estimated using BioGeoBEARS. The DIVALIKE model (in bold) was selected for downstream analyses based on ΔAIC scores.

| Model | LnL | Number of parameters | *d* | *e* | AIC | ΔAIC |
| --- | --- | --- | --- | --- | --- | --- |
| DEC | -143.8 | 2 | 0.06 | 0.04 | 291.8 | 1.8 |
| **DIVALIKE** | -142.9 | 2 | 0.07 | 0.03 | 290 | 0 |
| BAYAREALIKE | -157.4 | 2 | 0.07 | 0.13 | 319.1 | 29.1 |

Ancestral range reconstruction based on the DIVALIKE model estimated that the most probable ancestral range for the AAG is a combination of the monsoonal tropics + arid zone (*P* = 0.49) followed by the monsoonal tropics (0.29), and other state combinations (0.32) (Fig.). The two monophyletic crown groups AG2 and DRE were inferred to have different ancestral ranges: the latter most likely originated in the mesic south-western zone + arid zone (0.46) followed by the arid zone (0.39); while the former probably in north-northeastern Australia (monsoonal tropics + arid zone; 0.61) (**Fig. 2**).

**Figure 2:**
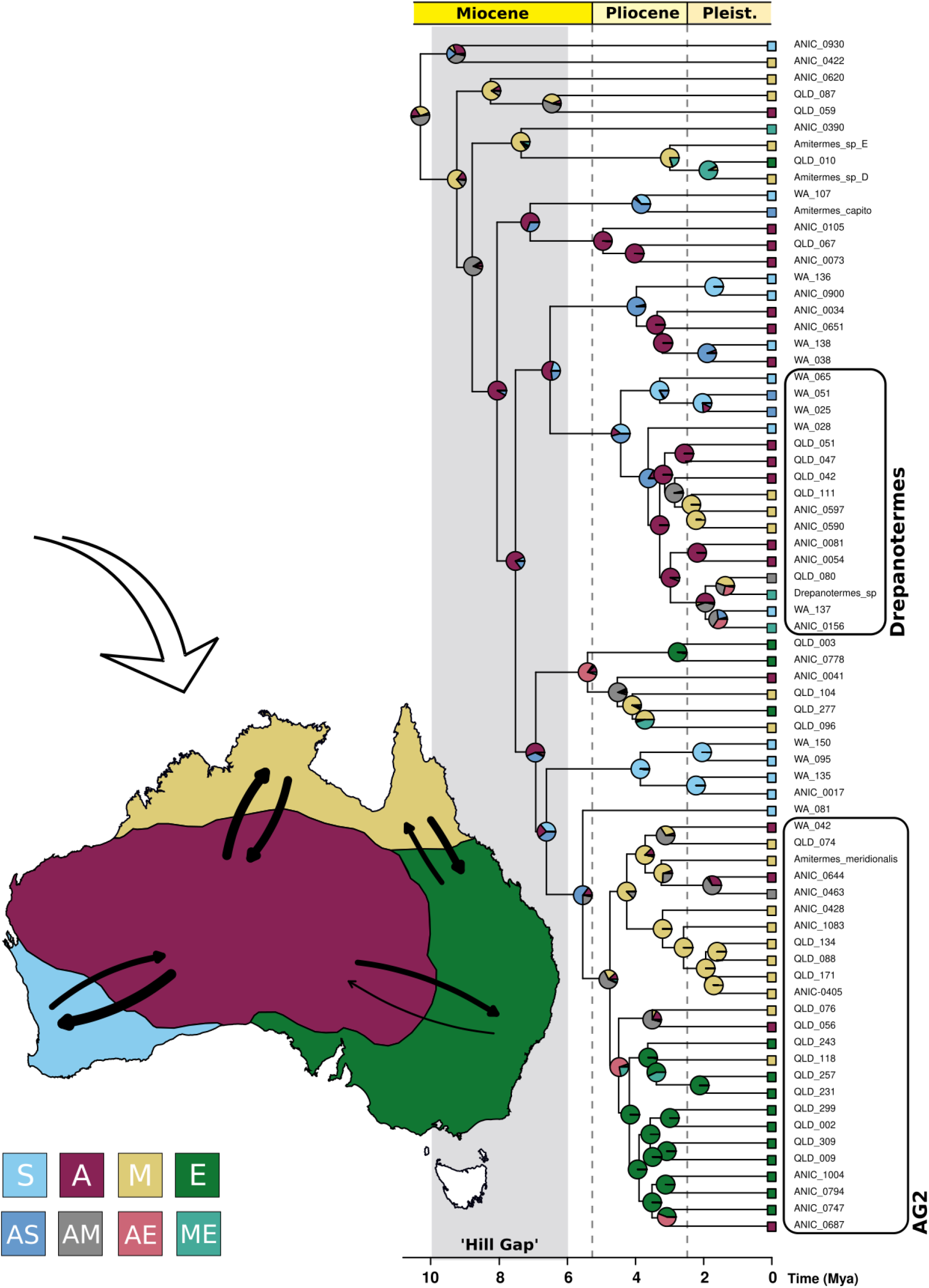
Ancestral range reconstruction of the AAG based on the DIVALIKE model. Relative probabilities of ancestral areas are shown in pie charts at nodes. Colored squares identify biomes: S, mesic south-western zone; A, arid zone; M, monsoonal tropics; E, mesic eastern zone. Combinations of biomes (*e.g*. AS, arid zone + mesic south-western zone) are also indicated in colored squares, but not shown on the map. The “Hill Gap” is shaded in grey. Black arrows indicate direction and frequency of dispersal events between biomes and line thickness indicates the number of event counts summarized with biogeographical stochastic mapping (BSM) (see **Tab. S4**). The white arrow indicates the putative arrival direction of the ancestor of the AAG (see Discussion). AAG taxa collected in this study are given with sample ID (see **Tab. S1**).

The BSM results indicate a complex biogeographical history, driven largely by within-area speciation and dispersal events (46.2% and 34.6% of the total number of events) rather than vicariance events (19.2%; **Tab. 2**). The highest number of dispersals occurred from the arid zone to the mesic south-western zone (~9 of 38 total estimated events), next highest was movements from the arid zone to the monsoonal tropics (~7 of 38) and from the monsoonal tropics to the arid zone (~6 of 38) (**Tab. S4**). More than half of all estimated dispersal events (52.1%) started in the arid zone, with the next common source being the monsoonal tropics (28.5%) (**Tab. S4** and **Fig. 2**). Most of these movements were directed towards the arid zone (28.1%), followed closely by the monsoonal tropics and the mesic south-western zone (26.6% and 24.6%, respectively) (**Tab. S4** and **Fig. 2**).

**Table 2:** Summary counts of 100 BSMs based on the DIVALIKE model using BioGeoBEARS. The estimated number of events for the different types are given in mean numbers with standard deviations (SD) and percentages.

|  | Range expansion | Within-area speciation | Vicariance | Total number of events |
| --- | --- | --- | --- | --- |
| mean | 37.51 | 50.15 | 20.85 | 108.5 |
| SD | ±1.7 | ±1.91 | ±1.91 | ±1.7 |
| % of total events | 34.6 | 46.2 | 19.2 | 100 |

### Model and rates of diversification

The LTT plot suggests two pulses of increased diversification followed by declines: (1) a first slight increase occurring ~9-6.5 Mya with a decline ~6.5-5.5 Mya; and (2) a subsequent abrupt, steep incline (~4.5-3 Mya) that exceeds the 95% confidence interval followed by a slowdown to the present day. The MCCR test showed a significantly negative γ statistic (−2.36, p = 9.999e-05) indicating non-constant species accumulation over time, as shown by the LTT plot (**Fig. 3**).

**Figure 3:**
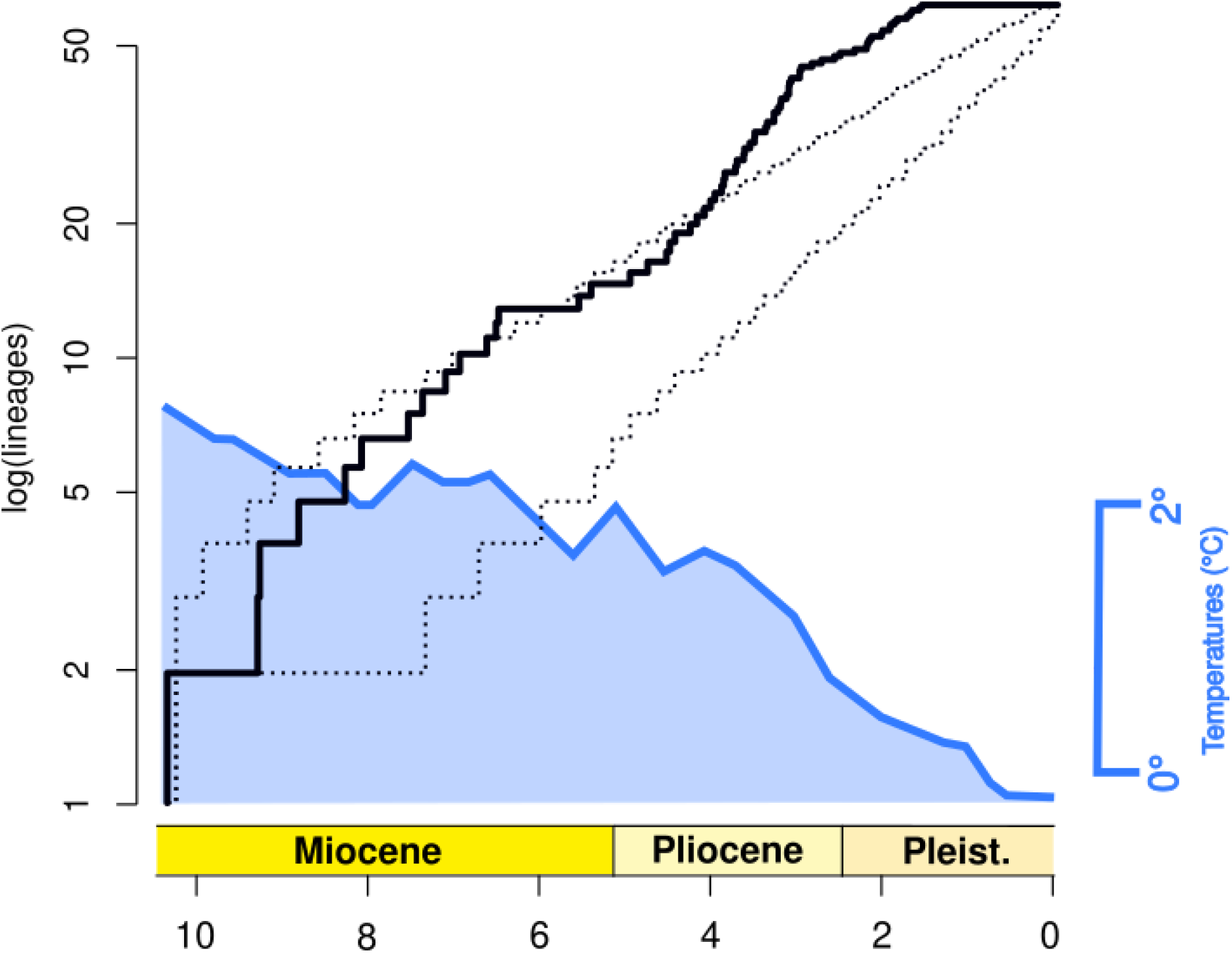
The LTT curve (black line), plotted against deep ocean temperatures (blue; modified from Zachos *et al*., 2001), shows two pulses of rapid diversification, which fall outside the 95% confidence interval generated by simulated LTTs (dashed lines).

The diversity-dependent linear model (DDL+E) was the best-fitting estimate of evolutionary diversification within the AAG (**Tab. 3**), based on four candidate models used in the maximum-likelihood diversification analyses. This supports again that a non-constant mode of diversification has shaped the evolutionary trajectory of the AAG. The same holds true when we assumed that only 48% or 29% of the ‘true’ diversity of the group was sampled in the phylogeny (**Tab. 3**), indicating that the results are robust against potential missing species.

**Table 3:** Alternative evolutionary models were fitted to the diversification history of the AAG. Included models are Yule (pure-birth model), CR (constant-rate birth death model), DDL+E (density-dependent linear diversification model), and DDX+E (density-dependent exponential diversification mode). The best-fit of models are based on ΔAICc scores.

| Model | logL | $\lambda$ | $\mu$ | AICc | $\Delta\text{AICc}$ |
| --- | --- | --- | --- | --- | --- |
| **72/100** |  |  |  |  |  |
| Yule | -167 | 0.29 | 0 | 336 | 27.7 |
| CR | -167 | 0.29 | 0 | 338.1 | 29.8 |
| DDL+E | -151 | 0.74 | 5.9e-05 | 308.3 | 0 |
| DDX+E | -179.5 | 0.57 | 0.35 | 365.3 | 57 |
| **72/150** |  |  |  |  |  |
| Yule | -162.8 | 0.36 | 0 | 327.6 | 17.5 |
| CR | -162.8 | 0.36 | 0 | 329.7 | 19.6 |
| DDL+E | -151.9 | 0.74 | 3.8e-06 | 310.1 | 0 |
| DDX+E | -161.9 | 1.52 | 0.12 | 330.1 | 20 |
| **72/250** |  |  |  |  |  |
| Yule | -158.4 | 0.45 | 0 | 318.78 | 0.98 |
| CR | -158.4 | 0.45 | 0 | 320.9 | 3.1 |
| DDL+E | -155.7 | 1.01 | 0.16 | 317.8 | 0 |
| DDX+E | -158.6 | 1.15 | 0.04 | 323.52 | 5.72 |

BAMM analyses showed no major shifts in the rate of diversification within the AAG (**Tab. S5** and **Fig. S4**), irrespective of the prior value that governs the number of rate shifts. The best-fitting run with the highest probability was fixed at 0.1 (**Tab. S5**). Consistent with a pattern of rapid diversification, the rate of diversification decreased markedly over time (**Fig. S4**), while the extinction rate remained near zero (mean extinction rate of ~0.05; **Fig. S4**)

## Discussion

The AAG is monophyletic and sister to *A. dentatus* from Southeast Asia. This indicates a single arrival event on the Australian continent through the Southeast Asian Archipelago about 11-10 Mya. This northern route is thought to have facilitated the dispersal of many insect species (Condamine *et al*., 2013, Matos-Maraví *et al*., 2018), after the Southeast Asian and Australian Plates collided 25-20 Mya (Hall, 2002). Our findings are in agreement with earlier results (Bourguignon *et al*., 2017); the missing lineages *Ahamitermes*, *Incolitermes*, and *Invasitermes*, are endemic inquilines in the nests of other termites (six species in total; Gay, 1955; Calaby, 1956; Abensperg-Traun & Perry, 1998) and share derived traits such as the near or complete loss of soldiers (Gay, 1968) and highly specialised mandibles (Miller, 1984). This suggests that they diversified *in situ* on the continent, similar to AG2 and *Drepanotermes*. In our PCoA, AG2 and *Drepanotermes* were clearly separated from AG1 and recovered in all phylogenetic analyses with high nodal support. They are nested among lineages of AG1, rendering *Amitermes*, as currently described, paraphyletic, which confirms the long-standing notion that these genera are derived from *Amitermes* (Hill, 1942; Watson, 1982; Miller, 1984).

Our reconstruction of the biogeographical history indicates that the ancestral range of the AAG is a combination of the monsoonal tropics + arid zone. This does not necessarily conflict with our phylogenetic reconstructions, but simply reflects the likely geographic distribution of the most recent common ancestor of extant AAG species. Irrespective of where the group originated, the available evidence (including deep nodes in our phylogeny) indicates that Australia’s monsoon region was the starting point for the radiation of the AAG across the continent. Accordingly, early lineages that speciated *in situ* may have been preadapted to seasonal climates, as their Southeast Asian ancestor evolved under similar climatic conditions (Bowman *et al*., 2010), allowing them to quickly expand their distributions in the monsoonal tropics.

Our estimate of ancestral ranges using the best-fitting model DIVALIKE suggests around 37 dispersal events during the radiation of the AAG, with 31 events occurring from or to the arid zone. While this may seem to be a very high number considering termites’ generally poor dispersal ability (Eggleton, 2000), our reconstructions of ancestral ranges and inference of biological processes (*i.e*., within-area speciation, range expansion, and vicariance) need to be considered in the context of gradual biome turnover during the last 15 million years in Australia. Early in the AAG radiation, environmental and climatic conditions were much different than today. For example, the arid zone as we know it, with iconic sand desert landscapes, is no older than one million years (Fujioka & Chappell, 2010) and formed in response to increasing aridification in the Miocene and Pleistocene (Byrne *et al*., 2008; 2018). Even today’s stony deserts did not develop until the end of the Pliocene (~3-2 Mya; Fujioka *et al*., 2005), while central Australia seems to have been covered by open woodlands and gallery forests at least since the late Miocene (Mao & Retallack, 2019; but see also Travouillon *et al*., 2009). This means the initial radiation of the AAG during the late Miocene occurred before major biogeographical barriers, e.g. the Great Sandy Desert in Western Australia, developed. Such a relatively benign habitat likely fostered successful expansion into or through the evolving arid zone and frequent dispersal into other biomes. Open sclerophyllous forests, including *Acacia* and *Eucalyptus*, have been widespread since the Paleogene (summarized in Crisp & Cook, 2013); these plants are primary food sources of AAG taxa in semi-arid and arid regions of Australia (Andersen & Jacklyn, 1993; Abensperg-Traun *et al*., 1995) and their widespread distribution certainly also contributed to successful range expansion of these termites.

As expected, within-area speciation is the most frequent type of event recovered in the BSM analysis. This reflects the large size of the regions and the associated predominance of lineages endemic to individual biomes. Such geographic restriction is in contrast to species occurrence records for some AAG taxa in our study, however high inter- and intraspecific variability made taxonomic reconciliation with historical morphology-based records impracticable. Nonetheless, shallow Plio-Pleistocene divergences in our phylogeny, particularly seen in AG2, tend to occur in close geographic proximity, likely reflecting phylogeographic structuring of species in response to habitat heterogeneity caused by intensifying aridification (Fujioka & Chappell, 2010).

The lowest number of events was inferred for vicariance, occurring about half as often as dispersal events. Biogeographic barriers have played an important role in shaping the present-day distribution of the continent’s plant and animal species (Crisp & Cook, 2007; Owen *et al*., 2017; Harms *et al*., 2019). They are mainly attributed to increasing aridity over the last 20 million years, particularly in central Australia (Cracraft, 1982), and to dramatic sea level fluctuations during Pleistocene interglacial periods (Zachos *et al*., 2001). We observed a strong correspondence between the frequency of vicariance events in the Plio-Pleistocene and the diversification of arid lineages from mesic and tropical ancestors, suggesting that the formation of potential vicariance barriers resulted in repeated allopatric speciation (Cracraft, 1982). This is consistent with complex diversification dynamics shown in other organisms (summarized in Byrne *et al*., 2018). However, we also observed diversification in the opposite direction in which mesic and tropical species diverged from arid ancestors, for example in *Drepanotermes*. Such a pattern has been recently shown in spinifex grasses (Toon *et al*., 2015), commonly harvested by *Drepanotermes* spp., which might indicate that the latter co-diversified with the former.

The LTT plot suggests that two pulses of rapid diversification occurred during the evolution of the AAG, corresponding to the initial radiation of early lineages (belonging to AG1) and to the radiation of the two derived groups, AG2 and *Drepanotermes*. Several factors may have separately or jointly influenced the evolutionary trajectory of the AAG, including these two surges of diversification. We will consider four possible factors in detail: (1) extinction, (2) predation, (3) key innovations, and (4) availability of niche space.

First, we consider it unlikely that an increase in extinction is responsible for the observed pattern, as the AAG diversified in the near absence of extinction. Legendre and Condamine (2018) showed that extinction rates are in general exceptionally low in termites compared to other dictyoptera, which is related to their eusocial lifestyle. Nonetheless, there are long naked branches in our phylogeny indicative of extinction events, and the current distribution of sister species in opposite corners of Australia suggests the occurrence of repeated range expansions and contractions coupled with extinction in the past. Yet there is no evidence that a mass extinction event took place that could have spurred rapid diversification (Losos, 2010). Instead, extinction seems to have been rare and sporadic within the AAG.

Second, although termites are consumed by many different animals including mammals, ants, and lizards (Holt, 1990; Abensperg-Traun & De Boer, 1992; dos Reis *et al*., 2012), predator-prey interactions are poorly understood (Noirot & Darlington, 2000). However, most termite-eating animals are found in the semi-arid and arid zones of Australia, and in those regions, termites make up the greatest proportions of their diets (Abensperg-Traun, 1994; Palmer, 2010). Termite-eating lizards, in particular, are abundant in the arid zone (Morton & James 1988), and there seems to be a positive link between lizard and termite richness in central Australia (Pianka, 1981; Colli *et al*., 2007). While *Amitermes* soldiers have been described as generally “the embodiment of cowardice and uselessness” (Hill, 1922), *Drepanotermes* soldiers are moderately large, produce copious amounts of defensive secretions, and have long sickle-shaped mandibles. They are numerous and can be highly effective against predatory ants (*e.g. Iridomyrmex*: Greenslade, 1970; for discussion of temporary nest occupation see Holt, 1990) as well as other insects and spiders (Hill, 1922) and vertebrates (echidnas: Abensperg-Traun & De Boer, 1992; lizards: Hill, 1922). This suggests a response to predation pressure during arid-zone evolution. Although it is very unlikely that predation alone can explain the macroevolutionary dynamics that shaped the diversification of the AAG, its impact should not be underestimated (see Nosil & Crespi, 2006; Hautmann, 2020).

Third, key innovations have been frequently invoked to explain high levels of species diversity (Yoder *et al*., 2010), and in a broad sense, any trait that promotes increasing diversification can qualify (Rabosky, 2017). Strikingly, the majority of mound-building species either belong to AG2 (*e.g. A. meriodionalis*) or *Drepanotermes* (*e.g. D. tamminensis*), which diversified rapidly in the Plio-Pleistocene. This may indicate a causal link between the evolution of mound-building and the second diversification surge in the early Pliocene. However, BAMM inferences showed no rate shift(s) associated with the acquisition of this trait; indeed no rate shifts at all were detected by BAMM. Regardless of whether mound-building is a key innovation or a subsequent adaptive change, mound-building species have become ecologically dominant in mesic and tropical regions (Andersen & Jacklyn, 1993; Abensperg-Traun & Perry, 1998), indicating an important role in the rapid diversification of AG2 and *Drepanotermes*.

Fourth, the two bursts in diversification rate coincide with major environmental and climatic changes in the late Miocene and early Pliocene. The contraction of rainforests and expansion of sclerophyllous vegetation in the late Miocene and the brief return of warm and wet conditions in the early Pliocene, including the re-expansion of rainforests (Byrne et al., 2008), represent an unparalleled source of ecological opportunity. Such a scenario is known to trigger rapid diversification (Losos, 2010). It is mainly invoked in island systems where resources are abundant and interspecific competition low (Gillespie, 2016), as opposed to continents with complex and competitive habitats (Rabosky & Lovette, 2008). However, Pincheira-Donoso *et al*. (2015) has shown that dramatic environmental changes, brought about by the Andean uplift ~25 Mya, facilitated the rapid continental radiation of *Liolaemus* lizards. The intensifying aridification that caused gradual biome turnover on the Australian continent in the last 20 million years may have created similar ecological “windows of opportunity” that closed as niches filled. This is supported by recovering the DDL+E model as the best-fitting estimate of evolutionary diversification within the AAG, implying a declining rate of diversification due to species accumulation and niche saturation over time.

Of course, incomplete taxon sampling can also contribute to patterns of decreasing diversification (Cusimano & Renner, 2010), such as the Pleistocene plateau in the LTT plot. We would expect this apparent rate change to be less abrupt with greater representation from the central deserts and NW Western Australia. In addition, *Amitermes* includes many soil-dwelling (and possibly soil-feeding) species, which are more often under-sampled in phylogenetic studies (Chouvenc *et al*., 2021). Despite these limitations, the large number of recent (< 2 Mya) splits reflect ongoing diversification. This can also be observed in the closely related species and species complexes of the AAG (*e.g. Drepanotermes perniger*; Watson & Perry, 1981), often in sympatric associations, and the high degree of endemism (Gay, 1968; Watson & Gay, 1991; Watson & Perry, 1981; Abensperg-Traun & Perry, 1998).

Our results indicate that the AAG has an immense potential to adapt to changing climatic conditions. Because the activity of termites is thought to increase the resistance of semi-/arid environments under the prospect of future climate change (Bonachela *et al*., 2015), the AAG could play an important role in maintaining Australia’s ecosystems in the face of human-mediated climate change. Current climate models predict a change over the next 50 years in Australia nearly as great as that of the last 10 million years with mean annual temperature increases of 1-6°C (Hughes, 2003). Future studies will show whether the AAG can adapt as quickly as it has in the past to keep up with the current pace of global climate change.

### Conclusion

This study illuminates the evolutionary history of the most speciose termite group in Australia and is one of the few biogeographical studies with a continent-wide focus. Consistent with dispersal patterns in other insects (Yeates & Cassis, 2017), the group’s ancestor arrived in Australia via a northern route from Southeast Asia. Despite being poor dispersers, early lineages were apparently able to expand their range quickly under favorable conditions in the late Miocene. The progressive aridification of the Australian continent and expansion of the arid zone, especially in the last 4 million years, has shaped the evolutionary trajectory of many, if not all, of the AAG lineages and continues to shape them. Multiple lines of evidence suggest that two pulses of rapid diversification occurred during the evolution of the AAG coincident with past global climate change. There are many studies showing that the intensifying aridification of the Australian continent triggered rapid diversification (summarized in Byrne *et al*., 2018), and this is the first study to do so for Australian termites. Congruent with diversity-dependent patterns, species accumulation declined likely due to progressive niche saturation. However, other factors such as key innovations or predation certainly play(ed) an important role in the rapid diversification of the AAG, however it remains to be seen to what extent.

Additional taxon sampling in underrepresented semi-/arid regions is necessary to answer open questions related to migration patterns during initial expansion across Australia and later diversification within the arid zone. This and other studies of the Australian fauna demonstrate the resilience of termites against naturally occurring environmental change. In this case, the aridification of the Australian continent was not an evolutionary dead end for the AAG but rather the impetus for adaptation and diversification.

## Author contributions

B.H. and T.R.H. designed research; B.H. and T.R.H. collected samples and determined species. B.H. and L.S. performed wet lab work. B.H. and A.B. analyzed data with input from T.R.H. B.H., A.B., S.S., and T.R.H. contributed to data interpretation and analyses. B.H. and T.R.H. wrote the paper with input from all authors.

## Acknowledgements

We like to thank Stephen Cameron for providing termite samples and we are grateful to Fabien Condamine for helpful insights into diversification analysis.

## Data availability

Mitochondrial genome sequences collected in this study will be deposited in GenBank.

