## Supplemental Material for "Rapid diversification of the Australian *Amitermes* group during late Cenozoic climate change"

**Long-range PCR**

Primer sequences were designed with Primer3 version 2.3.7 (Untergasser *et al.*, 2012) implemented in Geneious Prime. We used the available mitochondrial genome sequences of Australian *Amitermes* to design two pairs of primers spanning (a) ~8 kb and (b) ~10 kb fragments of the mitochondrial genome to ensure sufficient overlap between fragments. We used following forward and reverse primers: (a) nad1_ter_1 (ATCAAARGGWGTHCGATTMGTYTC) and cox2_ter_2 (TTTGCYCCRCARATYTCWGARCATTG); and (b) cox2_ter_3 (TGGCAGATAAGTGCRBTGGATTTAAG) and 16s_ter_4 (GAAGGGCCGCGGTATTTTGACC). The PrimeStar GXL polymerase (Takara Bio Europe) was used to amplify both fragments in a touchdown PCR procedure. The first 20 iterations were run as follows: the denaturing temperature was set to 98 °C (30 s), the initial annealing temperature of 60 °C (15 s) was gradually decreased to 50 °C with a step of 0.5 °C per cycle, and the elongation temperature was set to 62 °C (10 min). This was followed by 10 iterations of 98 °C (10 s), denaturing temperature; 50 °C (15 s), annealing temperature; and 62 °C (10 min), elongation temperature.

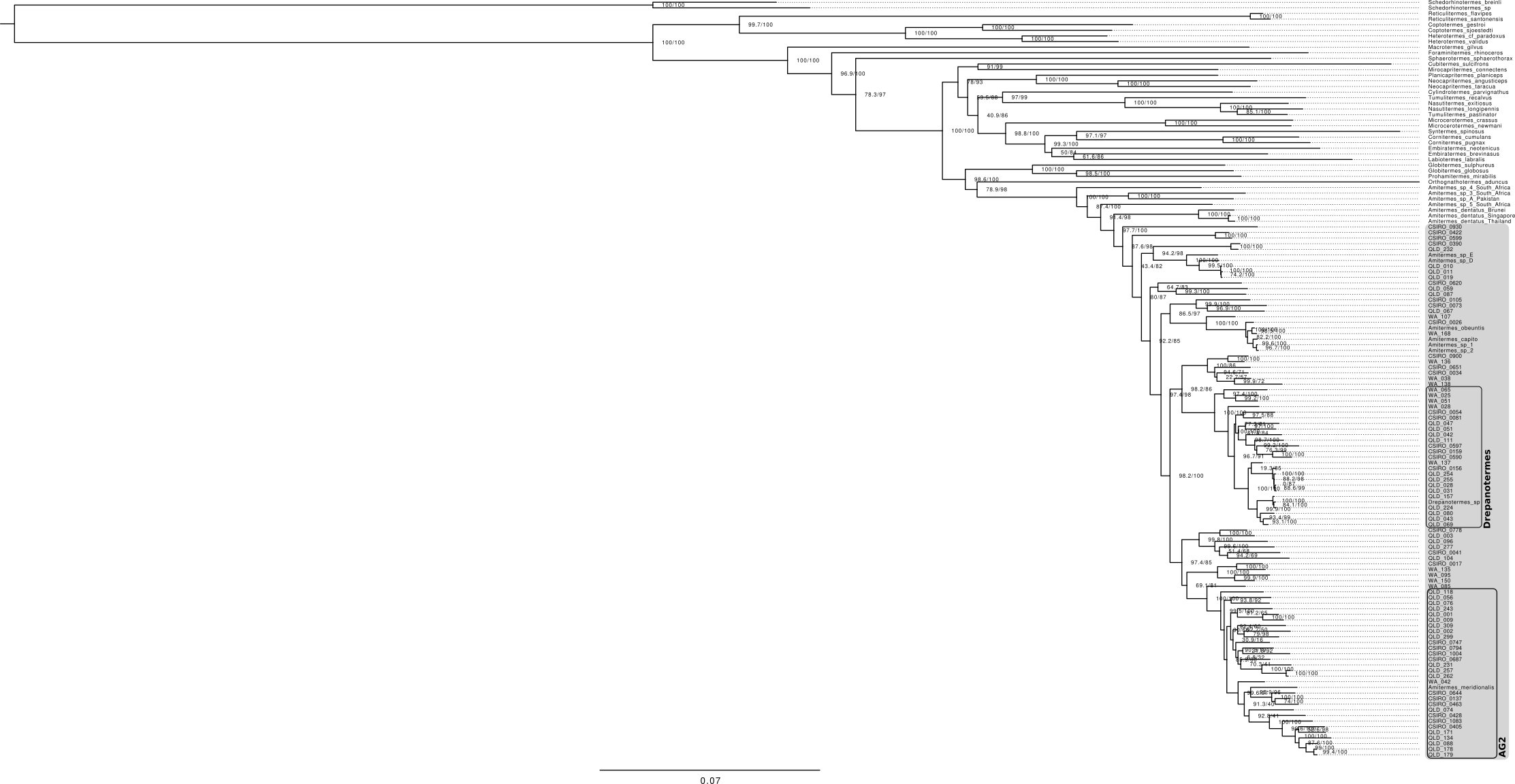

**Supplemental Figure 1**: Maximum likelihood tree inferred with IQ-TREE ver. 2.0.6. The AAG is indicated by the grey box, with *Drepanotermes* and AG2 in black boxes. Nodes are labelled with SH-aLRT/ufBS values. AAG taxa collected in this study are given with sample ID (see **Tab. S1**).

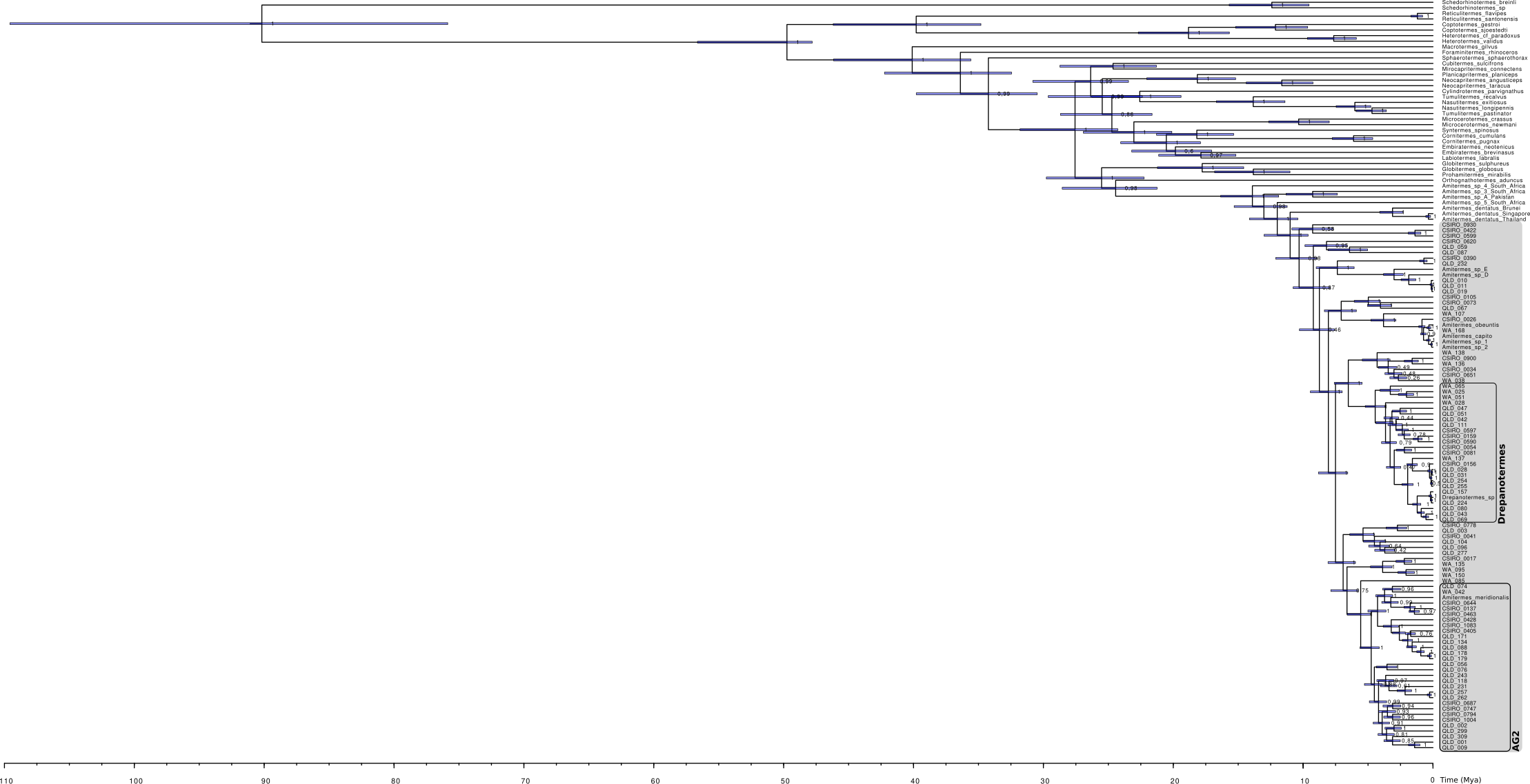

**Supplemental Figure 2:** Bayesian phylogenetic chronogram inferred with BEAST 2.6.1. The AAG is indicated by the grey box, with *Drepanotermes* and AG2 in black boxes. The scale bar is given in millions of years. Node bars represent the 95% credibility intervals of node-time estimates. Nodes are labelled with posterior probabilities. AAG taxa collected in this study are given with sample ID (see **Tab. S1**).

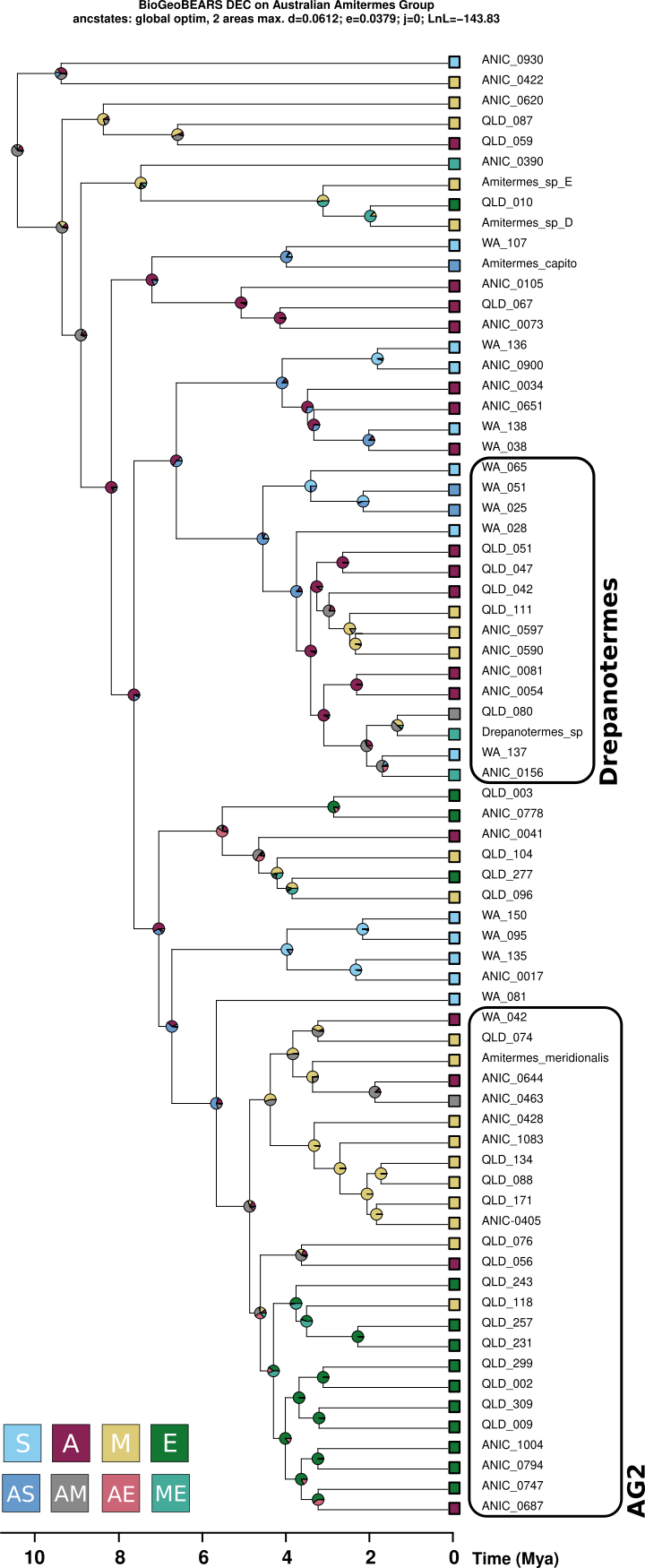

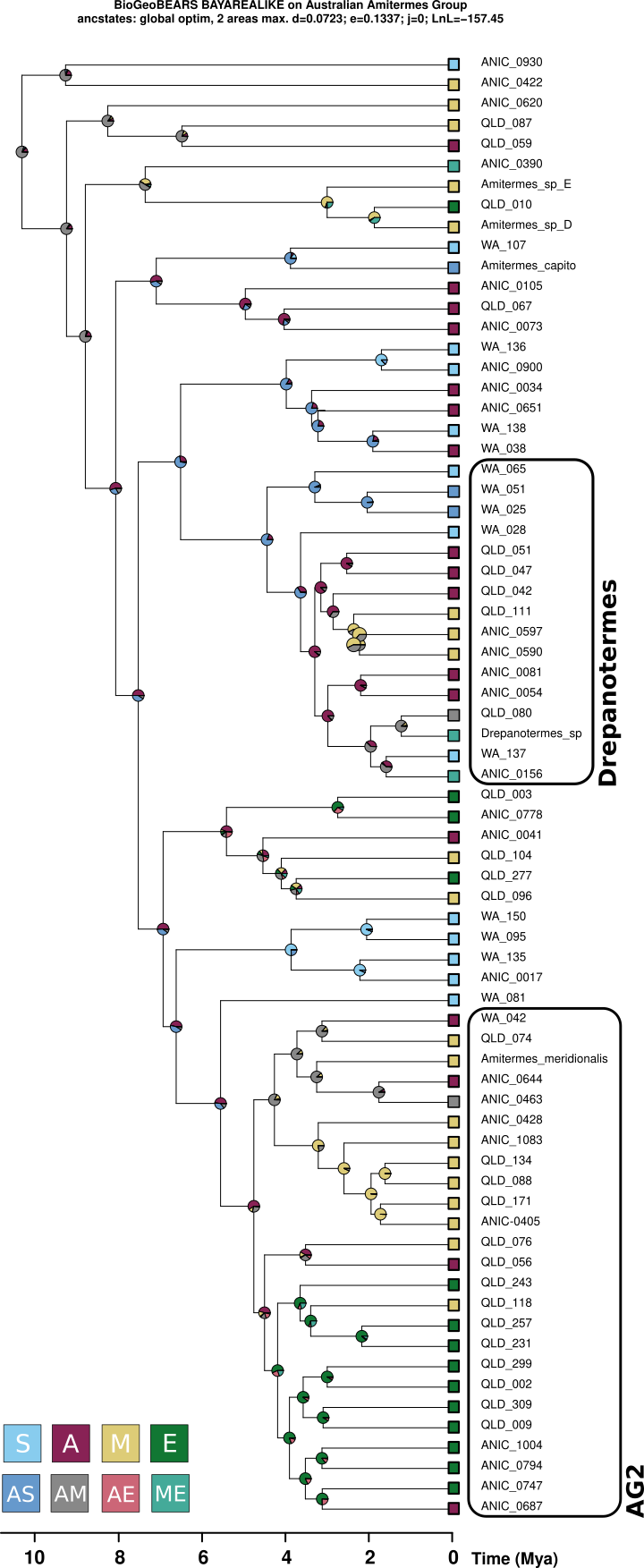

**Supplemental Figure 3:** Ancestral range reconstruction based on DEC and BAYAREALIKE models using BioGeoBEARS. Relative probabilities of ancestral areas are shown in pie charts at nodes. Colored squares identify biomes: S, mesic south-western zone; A, arid zone; M, monsoonal tropics; E, mesic eastern zone. Combinations of biomes (*e.g.* AS, arid zone + mesic south-western zone) are also indicated in colored squares, but not shown on the map. AAG taxa collected in this study are given with sample ID (see **Tab. S1**)

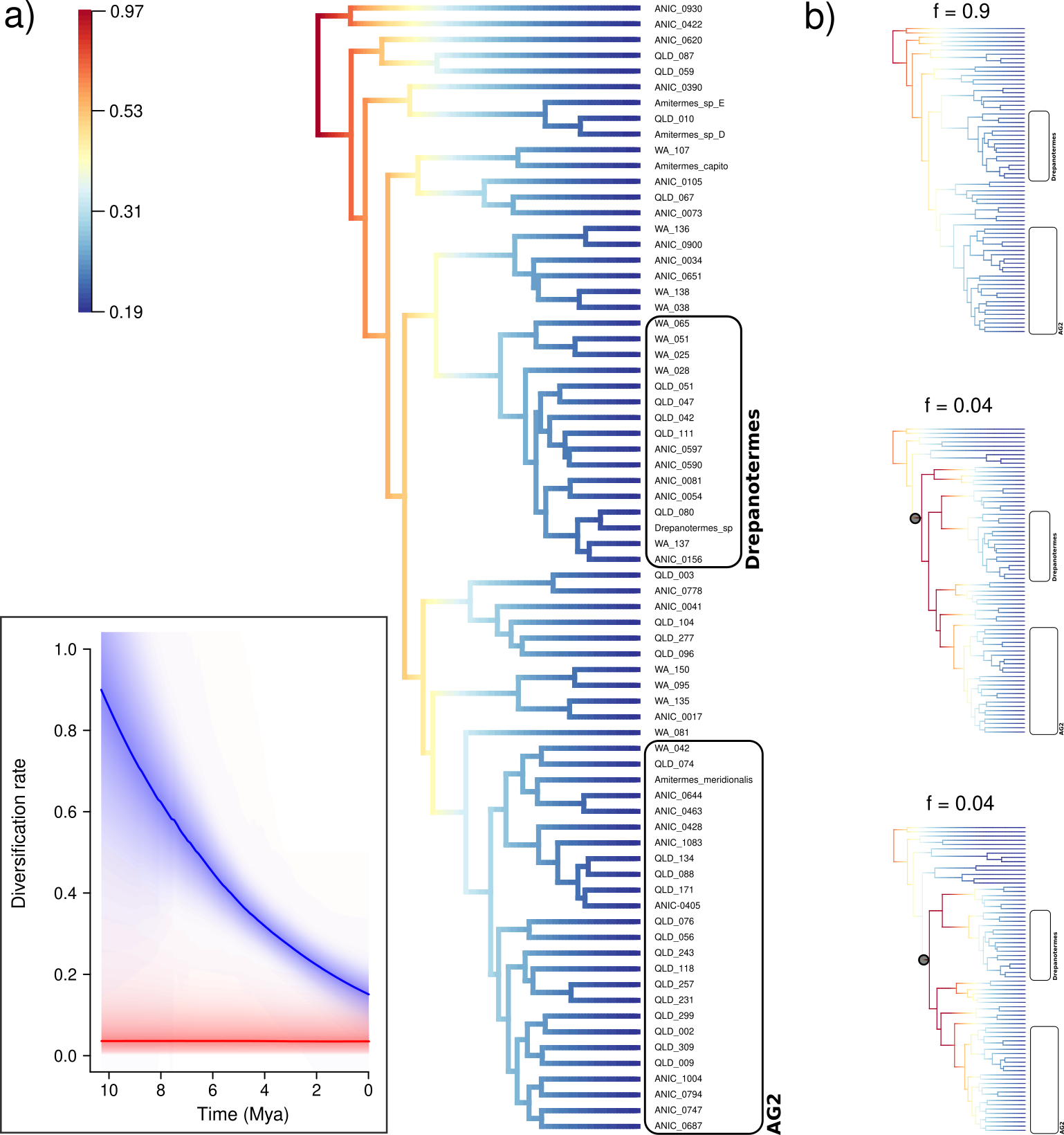

**Supplemental Figure 4:** Diversification pattern of the AAG inferred with BAMM 2.5.0. a) The best-fitting run showed no rate shift(s) in diversification. Warmer colors denote faster rates of diversification. Inset plot shows the mean diversification rate and mean extinction rate in blue and red lines, respectively. b) Plots depict the most likely distinct shift configurations for the best-fitting run (see **Tab. S5**), grey circles denote locations of rate shifts for each distinct shift configuration. AAG taxa collected in this study are given with sample ID (see **Tab. S1**).

**Supplemental Table 1:** Summary of sample information, including sample ID, collection locality, GenBank accession numbers, sequencing platform, and PCoA group.

| **Species** | **Sample ID** | **Collecting locality** | **Latitude** | **Longitude** | **Reference** | **Accession Number** | **Sequencing platform (read length)** | **PCoA group‡** |
| --- | --- | --- | --- | --- | --- | --- | --- | --- |
| *Amitermes capito*† |  | Western Australia, Australia |  |  | Bourguignon *et al.*, 2017 | KY224432 |  | AG1 |
| *Amitermes dentatus*† |  | Brunei, Asia |  |  | Bourguignon *et al.*, 2017 | KY224593 |  |  |
| *Amitermes dentatus*† |  | Singapore, Asia |  |  | Bourguignon *et al.*, 2017 | KY224549 |  |  |
| *Amitermes dentatus*† |  | Thailand, Asia |  |  | Bourguignon *et al.*, 2017 | KY224513 |  |  |
| *Amitermes meridionalis*† |  | Darwin, Northern Territory, Australia |  |  | Bourguignon *et al*., 2017 | KY224487 |  | AG2 |
| *Amitermes obeuntis*† |  | Western Australia, Australia |  |  | Bourguignon *et al*., 2017 | KY224650 |  | AG1 |
| *Amitermes* sp 1† |  | Western Australia, Australia |  |  | Bourguignon *et al.*, 2017 | KY224629 |  | AG1 |
| *Amitermes* sp 2† |  | Western Australia, Australia |  |  | Bourguignon *et al*., 2017 | KY224602 |  | AG1 |
| *Amitermes* sp 3† |  | South Africa, Africa |  |  | Bourguignon *et al*., 2017 | KY224581 |  |  |
| *Amitermes* sp 4† |  | South Africa, Africa |  |  | Bourguignon *et al.*, 2017 | KY224426 |  |  |
| *Amitermes* sp 5† |  | South Africa, Africa |  |  | Bourguignon *et al*., 2017 | KY224528 |  |  |
| *Amitermes* sp A† |  | Pakistan, Asia |  |  | Bourguignon *et al*., 2017 | KY224621 |  |  |
| *Amitermes* sp D† |  | Cairns, Queensland, Australia |  |  | Bourguignon *et al.*, 2017 | KY224695 |  | AG1 |
| *Amitermes* sp E† |  | Cairns, Queensland, Australia |  |  | Bourguignon *et al*., 2017 | KY224564 |  | AG1 |
| *Drepanotermes* sp† |  | Queensland, Australia |  |  | Cameron *et al*., 2012 | JX144938 |  | DRE |
| *Amitermes* sp | ANIC_0017 | Western Australia, Australia | -30.2 | 116 |  |  | MGISEQ-2000 (150 bp) | AG1 |
| *Amitermes* sp | ANIC_0026 | Western Australia, Australia | -29.7 | 116.2 |  |  | MGISEQ-2000 (150 bp) | AG1 |
| *Amitermes* sp | ANIC_0034 | Western Australia, Australia | -28.1 | 117.8 |  |  | MGISEQ-2000 (150 bp) | AG1 |
| *Amitermes* sp | ANIC_0041 | Western Australia, Australia | -27.4 | 117.9 |  |  | MGISEQ-2000 (150 bp) | AG1 |
| *Drepanotermes* sp | ANIC_0054 | Western Australia, Australia | -26 | 118.7 |  |  | MGISEQ-2000 (150 bp) | DRE |
| *Amitermes* sp | ANIC_0073 | Western Australia, Australia | -23.2 | 119.5 |  |  | MGISEQ-2000 (150 bp) | AG1 |
| *Drepanotermes* sp | ANIC_0081 | Western Australia, Australia | -21.7 | 118.8 |  |  | MGISEQ-2000 (150 bp) | DRE |
| *Amitermes* sp | ANIC_0105 | Western Australia, Australia | -19.8 | 121.1 |  |  | MGISEQ-2000 (150 bp) | AG1 |
| *Amitermes* sp | ANIC_0137 | Western Australia, Australia | -18.7 | 126.2 |  |  | MGISEQ-2000 (150 bp) | AG2 |
| *Drepanotermes* sp | ANIC_0156 | Western Australia, Australia | -15.8 | 128.8 |  |  | MGISEQ-2000 (150 bp) | DRE |
| *Drepanotermes* sp | ANIC_0159 | Northern Territory, Australia | -16.1 | 129.2 |  |  | MGISEQ-2000 (150 bp) | DRE |
| *Amitermes* sp | ANIC_0390 | Queensland, Australia | -17.6 | 145.4 |  |  | MGISEQ-2000 (150 bp) | AG1 |
| *Amitermes* sp | ANIC_0405 | Queensland, Australia | -17.9 | 144.9 |  |  | MGISEQ-2000 (150 bp) | AG2 |
| *Amitermes* sp | ANIC_0422 | Queensland, Australia | -18.3 | 143.4 |  |  | MGISEQ-2000 (150 bp) | AG1 |
| *Amitermes* sp | ANIC_0428 | Queensland, Australia | -18.2 | 142.9 |  |  | MGISEQ-2000 (150 bp) | AG2 |
| *Amitermes* sp | ANIC_0463 | Queensland, Australia | -17.9 | 139.3 |  |  | MGISEQ-2000 (150 bp) | AG2 |
| *Drepanotermes* sp | ANIC_0590 | Northern Territory, Australia | -14.7 | 132.7 |  |  | MGISEQ-2000 (150 bp) | DRE |
| *Drepanotermes* sp | ANIC_0597 | Northern Territory, Australia | -14.7 | 132.7 |  |  | MGISEQ-2000 (150 bp) | DRE |
| *Amitermes* sp | ANIC_0599 | Northern Territory, Australia | -15.3 | 133.1 |  |  | MGISEQ-2000 (150 bp) | AG1 |
| *Amitermes* sp | ANIC_0620 | Northern Territory, Australia | -16.9 | 133.4 |  |  | MGISEQ-2000 (150 bp) | AG1 |
| *Amitermes* sp | ANIC_0644 | Northern Territory, Australia | -21 | 134.2 |  |  | MGISEQ-2000 (150 bp) | AG2 |
| *Amitermes* sp | ANIC_0651 | Northern Territory, Australia | -23.1 | 133.7 |  |  | MGISEQ-2000 (150 bp) | AG1 |
| *Amitermes* sp | ANIC_0687 | Southern Australia, Australia | -28.1 | 135.8 |  |  | MGISEQ-2000 (150 bp) | AG2 |
| *Amitermes* sp | ANIC_0747 | New South Wales, Australia | -33.5 | 145.5 |  |  | MGISEQ-2000 (150 bp) | AG2 |
| *Amitermes* sp | ANIC_0778 | Queensland, Australia | -27.6 | 152.9 |  |  | MGISEQ-2000 (150 bp) | AG1 |
| *Amitermes* sp | ANIC_0794 | Queensland, Australia | -24.3 | 146.7 |  |  | MGISEQ-2000 (150 bp) | AG2 |
| *Amitermes* sp | ANIC_0900 | Western Australia, Australia | -28.7 | 115.2 |  |  | MGISEQ-2000 (150 bp) | AG1 |
| *Amitermes* sp | ANIC_0930 | Western Australia, Australia | -33.3 | 115.8 |  |  | MGISEQ-2000 (150 bp) | AG1 |
| *Amitermes* sp | ANIC_1004 | New South Wales, Australia | -34.4 | 147.6 |  |  | MGISEQ-2000 (150 bp) | AG2 |
| *Amitermes* sp | ANIC_1083 | Queensland, Australia | -12.6 | 143.4 |  |  | MGISEQ-2000 (150 bp) | AG2 |
| *Amitermes* sp | QLD_001 | Queensland, Australia | -26.5 | 151.8 |  |  | MGISEQ-2000 (150 bp) | AG2 |
| *Amitermes* sp | QLD_002 | Queensland, Australia | -26.5 | 151.8 |  |  | MGISEQ-2000 (150 bp) | AG2 |
| *Amitermes* sp | QLD_003 | Queensland, Australia | -25.6 | 151.6 |  |  | MGISEQ-2000 (150 bp) | AG1 |
| *Amitermes* sp | QLD_009 | Queensland, Australia | -23.7 | 149.6 |  |  | MGISEQ-2000 (150 bp) | AG2 |
| *Amitermes* sp | QLD_010 | Queensland, Australia | -23.7 | 149.6 |  |  | MGISEQ-2000 (150 bp) | AG1 |
| *Amitermes* sp | QLD_011 | Queensland, Australia | -23.7 | 149.6 |  |  | MGISEQ-2000 (150 bp) | AG1 |
| *Amitermes* sp | QLD_019 | Queensland, Australia | -23.6 | 149.2 |  |  | MGISEQ-2000 (150 bp) | AG1 |
| *Drepanotermes* sp | QLD_028 | Queensland, Australia | -23.5 | 148 |  |  | MGISEQ-2000 (150 bp) | DRE |
| *Drepanotermes* sp | QLD_031 | Queensland, Australia | -23.5 | 147.8 |  |  | MGISEQ-2000 (150 bp) | DRE |
| *Drepanotermes* sp | QLD_042 | Queensland, Australia | -22.3 | 143 |  |  | MGISEQ-2000 (150 bp) | DRE |
| *Drepanotermes* sp | QLD_043 | Queensland, Australia | -21.1 | 141.1 |  |  | MGISEQ-2000 (150 bp) | DRE |
| *Drepanotermes* sp | QLD_047 | Queensland, Australia | -21 | 140.9 |  |  | MGISEQ-2000 (150 bp) | DRE |
| *Drepanotermes* sp | QLD_051 | Queensland, Australia | -20.7 | 140.5 |  |  | MGISEQ-2000 (150 bp) | DRE |
| *Amitermes* sp | QLD_056 | Queensland, Australia | -20.8 | 140.2 |  |  | MGISEQ-2000 (150 bp) | AG2 |
| *Amitermes* sp | QLD_059 | Queensland, Australia | -20.8 | 140.2 |  |  | MGISEQ-2000 (150 bp) | AG1 |
| *Amitermes* sp | QLD_067 | Queensland, Australia | -19.9 | 140.2 |  |  | MGISEQ-2000 (150 bp) | AG1 |
| *Drepanotermes* sp | QLD_069 | Queensland, Australia | -19.9 | 140.2 |  |  | MGISEQ-2000 (150 bp) | DRE |
| *Amitermes* sp | QLD_074 | Queensland, Australia | -19.1 | 140.4 |  |  | MGISEQ-2000 (150 bp) | AG2 |
| *Amitermes* sp | QLD_076 | Queensland, Australia | -18.7 | 140.5 |  |  | MGISEQ-2000 (150 bp) | AG2 |
| *Drepanotermes* sp | QLD_080 | Queensland, Australia | -18.5 | 140.7 |  |  | MGISEQ-2000 (150 bp) | DRE |
| *Amitermes* sp | QLD_087 | Queensland, Australia | -17.7 | 141 |  |  | MGISEQ-2000 (150 bp) | AG1 |
| *Amitermes* sp | QLD_088 | Queensland, Australia | -18 | 141.4 |  |  | MGISEQ-2000 (150 bp) | AG2 |
| *Amitermes* sp | QLD_096 | Queensland, Australia | -18.2 | 142.3 |  |  | MGISEQ-2000 (150 bp) | AG1 |
| *Amitermes* sp | QLD_104 | Queensland, Australia | -18.2 | 142.9 |  |  | MGISEQ-2000 (150 bp) | AG1 |
| *Drepanotermes* sp | QLD_111 | Queensland, Australia | -18.1 | 144.4 |  |  | MGISEQ-2000 (150 bp) | DRE |
| *Amitermes* sp | QLD_118 | Queensland, Australia | -18 | 144.9 |  |  | MGISEQ-2000 (150 bp) | AG2 |
| *Amitermes* *laurensis* | QLD_134 | Queensland, Australia | -17.1 | 145.1 |  |  | MGISEQ-2000 (150 bp) | AG2 |
| *Drepanotermes* sp | QLD_157 | Queensland, Australia | -16.5 | 145.1 |  |  | MGISEQ-2000 (150 bp) | DRE |
| *Amitermes* sp | QLD_171 | Queensland, Australia | -15.8 | 144.7 |  |  | MGISEQ-2000 (150 bp) | AG2 |
| Amitermes sp | QLD_178 | Queensland, Australia | -15.7 | 144.6 |  |  | MGISEQ-2000 (150 bp) | AG2 |
| *Amitermes* sp | QLD_179 | Queensland, Australia | -15.6 | 144.5 |  |  | MGISEQ-2000 (150 bp) | AG2 |
| *Drepanotermes* sp | QLD_224 | Queensland, Australia | -19.9 | 146.6 |  |  | MGISEQ-2000 (150 bp) | DRE |
| *Amitermes* sp | QLD_231 | Queensland, Australia | -20.3 | 146.2 |  |  | MGISEQ-2000 (150 bp) | AG2 |
| *Amitermes* sp | QLD_232 | Queensland, Australia | -20.3 | 146.2 |  |  | MGISEQ-2000 (150 bp) | AG1 |
| *Amitermes* sp | QLD_243 | Queensland, Australia | -21 | 146.4 |  |  | MGISEQ-2000 (150 bp) | AG2 |
| *Drepanotermes* sp | QLD_254 | Queensland, Australia | -21.9 | 147 |  |  | MGISEQ-2000 (150 bp) | DRE |
| *Drepanotermes* sp | QLD_255 | Queensland, Australia | -22.8 | 147.6 |  |  | MGISEQ-2000 (150 bp) | DRE |
| *Amitermes* sp | QLD_257 | Queensland, Australia | -22.8 | 147.6 |  |  | MGISEQ-2000 (150 bp) | AG2 |
| *Amitermes* sp | QLD_262 | Queensland, Australia | -22.8 | 147.6 |  |  | MGISEQ-2000 (150 bp) | AG2 |
| *Amitermes* sp | QLD_277 | Queensland, Australia | -23.1 | 148 |  |  | MGISEQ-2000 (150 bp) | AG1 |
| *Amitermes* sp | QLD_299 | Queensland, Australia | -25.7 | 148.7 |  |  | MGISEQ-2000 (150 bp) | AG2 |
| *Amitermes* sp | QLD_309 | Queensland, Australia | -26.7 | 150.3 |  |  | MGISEQ-2000 (150 bp) | AG2 |
| *Drepanotermes gayi* | WA_025 | Western Australia, Australia | -30.3 | 116.7 |  |  | Illumina MiSeq (300 bp) | DRE |
| *Drepanotermes* sp | WA_028 | Western Australia, Australia | -30.3 | 116.7 |  |  | Illumina MiSeq (300 bp) | DRE |
| *Amitermes* *deplanatus* | WA_038 | Western Australia, Australia | -29.3 | 117.7 |  |  | Illumina MiSeq (300 bp) | AG1 |
| *Amitermes* sp | WA_042 | Western Australia, Australia | -29.3 | 117.7 |  |  | Illumina MiSeq (300 bp) | AG2 |
| *Drepanotermes clarki* | WA_051 | Western Australia, Australia | -27.9 | 117.9 |  |  | Illumina MiSeq (300 bp) | DRE |
| *Drepanotermes tamminensis* | WA_065 | Western Australia, Australia | -28.5 | 115.6 |  |  | Illumina MiSeq (300 bp) | DRE |
| *Amitermes* *heterognathus* | WA_085 | Western Australia, Australia | -29.5 | 115.4 |  |  | Illumina MiSeq (300 bp) | AG1 |
| *Amitermes* sp | WA_095 | Western Australia, Australia | -30 | 116.1 |  |  | Illumina MiSeq (300 bp) | AG1 |
| *Amitermes* sp | WA_107 | Western Australia, Australia | -31.7 | 116.5 |  |  | Illumina MiSeq (300 bp) | AG1 |
| *Amitermes* sp | WA_135 | Western Australia, Australia | -30.8 | 121.6 |  |  | Illumina MiSeq (300 bp) | AG1 |
| *Amitermes* sp | WA_136 | Western Australia, Australia | -31.3 | 119.7 |  |  | Illumina MiSeq (300 bp) | AG1 |
| *Drepanotermes* sp | WA_137 | Western Australia, Australia | -31.3 | 119.7 |  |  | Illumina MiSeq (300 bp) | DRE |
| *Amitermes* *conformis* | WA_138 | Western Australia, Australia | -32.4 | 118.4 |  |  | Illumina MiSeq (300 bp) | AG1 |
| *Amitermes* sp | WA_150 | Western Australia, Australia | -32.5 | 118.1 |  |  | Illumina MiSeq (300 bp) | AG1 |
| *Amitermes* *obeuntis* | WA_168 | Western Australia, Australia | -32.1 | 117.8 |  |  | Illumina MiSeq (300 bp) | AG1 |
| *Coptotermes gestroi* |  |  |  |  | Bourguignon *et al.*, 2016 | NC_030014 |  |  |
| *Coptotermes sjoestedti* |  |  |  |  | Bourguignon *et al.*, 2016 | NC_030020 |  |  |
| *Cornitermes cumulans* |  |  |  |  | Bourguignon *et al.*, 2017 | NC_034086 |  |  |
| *Cornitermes pugnax* |  |  |  |  | Bourguignon *et al.*, 2017 | NC_034055 |  |  |
| *Cubitermes sulcifrons* |  |  |  |  | Bourguignon *et al*., 2017 | NC_034109 |  |  |
| *Cylindrotermes parvignathus* |  |  |  |  | Bourguignon *et al.*, 2017 | KY224565 |  |  |
| *Embiratermes brevinasus* |  |  |  |  | Bourguignon *et al*., 2017 | NC_034101 |  |  |
| *Embiratermes neotenicus* |  |  |  |  | Bourguignon *et al*., 2017 | NC_034930 |  |  |
| *Foraminitermes rhinoceros* |  |  |  |  | Bourguignon *et al*., 2017 | NC_034116 |  |  |
| *Globitermes globosus* |  |  |  |  | Bourguignon *et al.*, 2017 | NC_034095 |  |  |
| *Globitermes sulphureus* |  |  |  |  | Bourguignon *et al*., 2017 | NC_034139 |  |  |
| *Heterotermes* cf *paradoxus* |  |  |  |  | Bourguignon *et al.,* 2016 | NC_030023 |  |  |
| *Heterotermes validus* |  |  |  |  | Bourguignon *et al*., 2016 | NC_030034 |  |  |
| *Labiotermes labralis* |  |  |  |  | Herve & Brune, 2017 | NC_034929 |  |  |
| *Macrotermes gilvus* |  |  |  |  | Bourguignon *et al*., 2017 | NC_034110 |  |  |
| *Microcerotermes crassus* |  |  |  |  | Bourguignon *et al.*, 2017 | NC_034036 |  |  |
| *Microcerotermes newmani* |  |  |  |  | Bourguignon *et al.*, 2017 | NC_034021 |  |  |
| *Nasutitermes exitiosus* |  |  |  |  | Bourguignon *et al.*, 2017 | NC_034115 |  |  |
| *Mirocapritermes connectens* |  |  |  |  | Bourguignon *et al.,* 2017 | NC_034085 |  |  |
| *Nasutitermes longipennis* |  |  |  |  | Bourguignon *et al.*, 2017 | NC_034060 |  |  |
| *Neocapritermes angusticeps* |  |  |  |  | Bourguignon *et al.*, 2017 | NC_034053 |  |  |
| *Neocapritermes taracua* |  |  |  |  | Dietrich & Brune, 2016 | NC_026116 |  |  |
| *Orthognathotermes aduncus* |  |  |  |  | Bourguignon *et al*., 2015 | KP026289 |  |  |
| *Planicapritermes planiceps* |  |  |  |  | Bourguignon *et al*., 2017 | NC_034090 |  |  |
| *Prohamitermes mirabilis* |  |  |  |  | Bourguignon *et al.*, 2017 | NC_034039 |  |  |
| *Reticulitermes flavipes* |  |  |  |  | Cameron & Whiting, 2007 | EF206314 |  |  |
| *Reticulitermes santonensis* |  |  |  |  | Cameron & Whiting, 2007 | EF206315 |  |  |
| *Schedorhinotermes breinli* |  |  |  |  | Cameron et *al.*, 2012 | NC_018126 |  |  |
| *Schedorhinotermes* sp |  |  |  |  | Wang *et al*., 2019 | MK246859 |  |  |
| *Sphaerotermes sphaerothorax* |  |  |  |  | Bourguignon *et al*., 2017 | NC_034103 |  |  |
| *Syntermes spinosus* |  |  |  |  | Bourguignon *et al.*, 2015 | KP026293 |  |  |
| *Tumulitermes pastinator* |  |  |  |  | Bourguignon *et al.*, 2017 | NC_034098 |  |  |
| *Tumulitermes recalvus* |  |  |  |  | Bourguignon *et al*., 2017 | NC_034051 |  |  |

†Sequences used for customized mitochondrial database in Kraken2.

‡According to PCoA analysis.

**Supplemental Table 2:** Fossils used as internal calibrations for divergence dating with BEAST 2.6.1.

| **Species** | **Minimum age constraint (Mya)** | **Calibration group** | **Soft maximum bound (97.5% probability)†** | **Note on maximum bond** |
| --- | --- | --- | --- | --- |
| *Nanotermes isaacae* | 47.8 | Termitidae + *Reticulitermes* + *Coptotermes* + *Heterotermes* | 93.5 | *Archeorhinotermes rossi*, first fossil of Rhinotermitidae |
| *Reticulitermes antiquus* | 33.9 | *Reticulitermes + Coptotermes + Heterotermes* | 93.5 |  |
| *Microcerotermes insularis* | 13.8 | *Microcerotermes* + Syntermitinae | 47.8 | *Nanotermes isaacae*, first fossil of Termitidae |
| *Amitermes lucidus* | 13.8 | *Amitermes + Globitermes + Orthognathotermes + Prohamitermes* | 47.8 |  |

†Fossil age estimates taken from the Paleobiology Database (www.paleobiodb.org; accessed 30 July 2020).

**Supplemental Table 3:** AAG taxa included in ancestral range reconstruction. Biogeographical distribution in the four major biomes given by presence (1) or absence (0). Biomes are abbreviated as follows: S, mesic south-western zone; A, arid zone; M, monsoonal tropics; E, mesic eastern zone. Sequences pruned from the time calibrated tree are given with sample ID (see Tab. S1) after their corresponding species.

| **Species** | **Sample code** | **Pruned sequences** | **S** | **A** | **M** | **E** |
| --- | --- | --- | --- | --- | --- | --- |
| *Amitermes capito* |  | *Amitermes* sp 1, *Amitermes* sp 2 , *Amitermes* *obeuntis*, WA_168, ANIC_0026 | 1 | 1 | 0 | 0 |
| *Amitermes meridionalis* |  |  | 0 | 0 | 1 | 0 |
| *Amitermes* sp D |  |  | 0 | 0 | 1 | 0 |
| *Amitermes* sp E |  |  | 0 | 0 | 1 | 0 |
| *Drepanotermes* sp |  | QLD_157, QLD_224 | 0 | 0 | 1 | 1 |
| *Amitermes* sp | ANIC_0017 |  | 1 | 0 | 0 | 0 |
| *Amitermes* sp | ANIC_0034 |  | 0 | 1 | 0 | 0 |
| *Amitermes* sp | ANIC_0041 |  | 0 | 1 | 0 | 0 |
| *Drepanotermes* sp | ANIC_0054 |  | 0 | 1 | 0 | 0 |
| *Amitermes* sp | ANIC_0073 |  | 0 | 1 | 0 | 0 |
| *Drepanotermes* sp | ANIC_0081 |  | 0 | 1 | 0 | 0 |
| *Amitermes* sp | ANIC_0105 |  | 0 | 1 | 0 | 0 |
| *Drepanotermes* sp | ANIC_0156 | QLD 028, QLD 031, QLD 254, QLD 255 | 0 | 0 | 1 | 1 |
| *Amitermes* sp | ANIC_0390 | QLD_232 | 0 | 0 | 1 | 1 |
| *Amitermes* sp | ANIC_0405 |  | 0 | 0 | 1 | 0 |
| *Amitermes* sp | ANIC_0422 | ANIC_0599 | 0 | 0 | 1 | 0 |
| *Amitermes* sp | ANIC_0428 |  | 0 | 0 | 1 | 0 |
| *Amitermes* sp | ANIC_0463 | ANIC_0137 | 0 | 1 | 1 | 0 |
| *Drepanotermes* sp | ANIC_0590 | ANIC_0159 | 0 | 0 | 1 | 0 |
| *Drepanotermes* sp | ANIC_0597 |  | 0 | 0 | 1 | 0 |
| *Amitermes* sp | ANIC_0620 |  | 0 | 0 | 1 | 0 |
| *Amitermes* sp | ANIC_0644 |  | 0 | 1 | 0 | 0 |
| *Amitermes* sp | ANIC_0651 |  | 0 | 1 | 0 | 0 |
| *Amitermes* sp | ANIC_0687 |  | 0 | 1 | 0 | 0 |
| *Amitermes* sp | ANIC_0747 |  | 0 | 0 | 0 | 1 |
| *Amitermes* sp | ANIC_0778 |  | 0 | 0 | 0 | 1 |
| *Amitermes* sp | ANIC_0794 |  | 0 | 0 | 0 | 1 |
| *Amitermes* sp | ANIC_0900 |  | 1 | 0 | 0 | 0 |
| *Amitermes* sp | ANIC_0930 |  | 1 | 0 | 0 | 0 |
| *Amitermes* sp | ANIC_1004 |  | 0 | 0 | 0 | 1 |
| *Amitermes* sp | ANIC_1083 |  | 0 | 0 | 1 | 0 |
| *Amitermes* sp | QLD_002 |  | 0 | 0 | 0 | 1 |
| *Amitermes* sp | QLD_003 |  | 0 | 0 | 0 | 1 |
| *Amitermes* sp | QLD_009 | QLD_001 | 0 | 0 | 0 | 1 |
| *Amitermes* sp | QLD_010 | QLD_011, QLD_019 | 0 | 0 | 0 | 1 |
| *Drepanotermes* sp | QLD_042 |  | 0 | 1 | 0 | 0 |
| *Drepanotermes* sp | QLD_047 |  | 0 | 1 | 0 | 0 |
| *Drepanotermes* sp | QLD_051 |  | 0 | 1 | 0 | 0 |
| *Amitermes* sp | QLD_056 |  | 0 | 1 | 0 | 0 |
| *Amitermes* sp | QLD_059 |  | 0 | 1 | 0 | 0 |
| *Amitermes* sp | QLD_067 |  | 0 | 1 | 0 | 0 |
| *Amitermes* sp | QLD_074 |  | 0 | 0 | 1 | 0 |
| *Amitermes* sp | QLD_076 |  | 0 | 0 | 1 | 0 |
| *Drepanotermes* sp | QLD_080 | QLD_043, QLD_069 | 0 | 1 | 1 | 0 |
| *Amitermes* sp | QLD_087 |  | 0 | 0 | 1 | 0 |
| *Amitermes* sp | QLD_088 | QLD_178, QLD_179 | 0 | 0 | 1 | 0 |
| *Amitermes* sp | QLD_096 |  | 0 | 0 | 1 | 0 |
| *Amitermes* sp | QLD_104 |  | 0 | 0 | 1 | 0 |
| *Drepanotermes* sp | QLD_111 |  | 0 | 0 | 1 | 0 |
| *Amitermes* sp | QLD_118 |  | 0 | 0 | 1 | 0 |
| *Amitermes* *laurensis* | QLD_134 |  | 0 | 0 | 1 | 0 |
| *Amitermes* sp | QLD_171 |  | 0 | 0 | 1 | 0 |
| *Amitermes* sp | QLD_231 |  | 0 | 0 | 0 | 1 |
| *Amitermes* sp | QLD_243 |  | 0 | 0 | 0 | 1 |
| *Amitermes* sp | QLD_257 | QLD_262 | 0 | 0 | 0 | 1 |
| *Amitermes* sp | QLD_277 |  | 0 | 0 | 0 | 1 |
| *Amitermes* sp | QLD_299 |  | 0 | 0 | 0 | 1 |
| *Amitermes* sp | QLD_309 |  | 0 | 0 | 0 | 1 |
| *Drepanotermes gayi* | WA_025 |  | 1 | 1 | 0 | 0 |
| *Drepanotermes* sp | WA_028 |  | 1 | 0 | 0 | 0 |
| *Amitermes* *deplanatus* | WA_038 |  | 0 | 1 | 0 | 0 |
| *Amitermes* sp | WA_042 |  | 0 | 1 | 0 | 0 |
| *Drepanotermes clarki* | WA_051 |  | 1 | 1 | 0 | 0 |
| *Drepanotermes tamminensis* | WA_065 |  | 1 | 0 | 0 | 0 |
| *Amitermes* *heterognathus* | WA_085 |  | 1 | 0 | 0 | 0 |
| *Amitermes* sp | WA_095 |  | 1 | 0 | 0 | 0 |
| *Amitermes* sp | WA_107 |  | 1 | 0 | 0 | 0 |
| *Amitermes* sp | WA_135 |  | 1 | 0 | 0 | 0 |
| *Amitermes* sp | WA_136 |  | 1 | 0 | 0 | 0 |
| *Drepanotermes* sp | WA_137 |  | 1 | 0 | 0 | 0 |
| *Amitermes* *conformis* | WA_138 |  | 1 | 0 | 0 | 0 |
| *Amitermes* sp | WA_150 |  | 1 | 0 | 0 | 0 |

**Supplemental Table 4:** Dispersal event counts estimated with biogeographical stochastic mapping (BSM), averaged across 100 BSMs. Rows indicate where the lineage dispersed from and columns where the lineage dispersed to. Standard deviations are given in parentheses. On the margins (in grey), the sum and percentages of events (in parentheses) involving each area are given, in which rows denote the starting point of dispersal, and columns the dispersal destination. Biomes are abbreviated as follows: S, mesic south-western zone; A, arid zone; M, monsoonal tropics; E, mesic eastern zone.

|  |  | **to** | | | |  |
| --- | --- | --- | --- | --- | --- | --- |
|  |  | **S** | **A** | **M** | **E** |  |
| **from** | **S** |  | 3.37 (±1.87) | 0 | 0 | 3.37 (9%) |
|  | **A** | 9.22 (±1.82) |  | 7.45 (±2.08) | 2.88 (±0.9) | 19.55 (52.1%) |
|  | **M** | 0 | 5.81 (±1.75) |  | 4.88 (±1.07) | 10.69 (28.5%) |
|  | **E** | 0 | 1.39 (±0.67) | 2.51 (±1.24) |  | 3.9 (10.4%) |
|  |  | 9.22 (24.6%) | 10.57 (28.1%) | 9.96 (26.6%) | 7.76 (20.7%) | 37.51 (100%) |

**Supplemental Table 5:** Summary of models compared across a range of Poisson rate priors using BAMM version 2.5.0 and BAMMtools v.2.1

|  |  | **Poisson rate prior** | | | | | | | | | |
| --- | --- | --- | --- | --- | --- | --- | --- | --- | --- | --- | --- |
|  |  | **0.1‡** | **0.2** | **0.3** | **0.4** | **0.5** | **0.6** | **0.7** | **0.8** | **0.9** | **1** |
| **ESS N shifts** | | 681.3 | 678.14 | 681.53 | 827.79 | 945.97 | 1093.51 | 890.57 | 945.57 | 1064.32 | 1628.1 |
| **ESS lnL** | | 721.81 | 657.35 | 569.34 | 702.81 | 795.11 | 868.05 | 649.8 | 691.94 | 735.25 | 1056.45 |
| **Posterior distribution per number of shifts** | **0** | **0.9‡** | 0.84 | 0.79 | 0.71 | 0.66 | 0.59 | 0.62 | 0.62 | 0.61 | 0.62 |
|  | **1†** | 0.04 | 0.06 | 0.08 | 0.1 | 0.12 | 0.15 | 0.14 | 0.14 | 0.14 | 0.14 |
|  | **1†** | 0.04 | 0.06 | 0.07 | 0.1 | 0.12 | 0.15 | 0.14 | 0.14 | 0.14 | 0.14 |
|  | **1†** | NA | NA | 0.03 | 0.04 | 0.05 | 0.06 | 0.06 | 0.06 | 0.06 | 0.06 |
|  | **1†** | NA | NA | NA | NA | NA | 0.05 | NA | NA | NA | NA |

ESS, effective sample size; lnL, likelihood scores; NA, not applicable.

†Core shift(s) can occur at different branches in the phylogenetic tree.

‡Selected model according to the highest posterior distribution per number of shifts.
